# Brain and grammar: revealing electrophysiological basic structures with competing statistical models

**DOI:** 10.1101/2024.02.06.579088

**Authors:** Andrea Cometa, Chiara Battaglini, Fiorenzo Artoni, Matteo Greco, Robert Frank, Claudia Repetto, Franco Bottoni, Stefano F Cappa, Silvestro Micera, Emiliano Ricciardi, Andrea Moro

**Affiliations:** MoMiLab, IMT School for Advanced Studies Lucca, Piazza S.Francesco, 19, 55100 Lucca, Italy; The BioRobotics Institute and Department of Excellence in Robotics and AI, Scuola Superiore Sant’Anna, Viale Rinaldo Piaggio 34, Pontedera 56025, Italy; Cognitive Neuroscience (ICoN) Center, University School for Advanced Studies IUSS, Piazza Vittoria 15, Pavia 27100, Italy; Neurolinguistics and Experimental Pragmatics (NEP) Lab, University School for Advanced Studies IUSS Pavia, Piazza della Vittoria 15, 27100 Pavia, Italy; Department of Clinical Neurosciences, Faculty of Medicine, University of Geneva, 1, rue Michel-Servet, 1211 Genéve, Switzerland; Department of Linguistics, Yale University, 370 Temple St, New Haven, CT 06511, Unites States; Department of Psychology, Università Cattolica del Sacro Cuore, Largo A. Gemelli 1, Milan 20123, Italy; Istituto Clinico Humanitas, IRCCS, Via Alessandro Manzoni 56, Rozzano 20089, Italy; IRCCS Mondino Foundation National Institute of Neurology, Via Mondino 2, Pavia 27100, Italy; Bertarelli Foundation Chair in Translational NeuroEngineering, Center for Neuroprosthetics and School of Engineering, Ecole Polytechnique Federale de Lausanne, Campus Biotech, Chemin des Mines 9, Geneva, GE CH 1202, Switzerland

## Abstract

Acoustic, lexical and syntactic information is simultaneously processed in the brain. Therefore, distinguishing the electrophysiological activity pertaining to these components requires complex and indirect strategies. Capitalizing on previous works which factor out acoustic information, we could concentrate on the lexical and syntactic contribution to language processing by testing competing statistical models. We exploited EEG recordings and compared different surprisal models selectively involving lexical information, part of speech or syntactic structures in various combinations. EEG responses were recorded in 32 participants during listening to affirmative active declarative sentences and compared the activation corresponding to basic syntactic structures, such as noun phrases vs verb phrases. Lexical and syntactic processing activates different frequency bands, different time windows and different networks. Moreover, surprisal models based on part of speech inventory only do not explain well the electrophysiological data, while those including syntactic information do. Finally, we confirm previous measures obtained with intracortical recordings independently supporting the original hypothesis addressed here in a robust way.

## Introduction

During sentence comprehension, syntactic information is crucially intertwined with acoustic information [1], [2], [3]. This makes it difficult to decouple neural syntactic processing from other types of language processing, particularly in electrophysiological studies [1], [2]. To address this problem, we previously designed a set of stimuli composed of sentences containing homophonous parts [4]. These homophonous parts have the same acoustic content but different syntactic structures, i.e., they could be either noun phrases (NPs – article + noun) or verb phrases (VPs – clitic (pronoun) + verb). This was made possible by exploiting three characteristics of the Italian language: (i) some definite articles are pronounced exactly like some object clitic pronouns (such as [la] written as *la*; it can be both “the - fem.sing.” or “her - fem.sing.”); (ii) the syntax of articles and clitic pronouns is very different: they both precede the noun/verb, but usually complements follow verbs. The operation of placing a pronoun before the verb is called *cliticization* [5]. The placement of the clitic pronoun before the verb has been taken to implicate a complex syntactic operation, which is absent in NPs. And (iii) the Italian lexicon contains several homophonous pairs of nouns and verbs, such as [ˈpɔrta] (written *porta*), which can either mean “door” or “brings”. Pairs of words such as [laˈpɔrta] (written as *la porta*) can thus be interpreted either as a noun phrase (“the door”) or a verb phrase (“brings her”) depending on the syntactic context (homophonous phrases - HPs).

In our previous works [4], [6], we compared the brain activity elicited by the processing of noun phrases or verb phrases, using stereo-electroencephalographic (SEEG) recordings. We found that the frequencies in the high-gamma band (150-300 Hz) were the main neural correlate of syntactic processing. We also observed a higher number of responsive contacts for VPs than for NPs, with the neural network supporting the processing of VPs being wider than the network processing NPs, and involving more cortical and subcortical areas, especially in the right (non-dominant) hemisphere.

A potential interpretation of these results comes from the notion of *surprisal*. Surprisal is defined as the negative log probability of a word in a context, which yields an inverse relationship between a word’s probability and its surprisal value [7]: the rarer a word is in a given context, the higher the surprisal. Surprisal is known to be positively correlated with brain activity [8].

Computing a surprisal value depends on the way in which a word’s probability is determined, i.e., what kind of language model is used. There are two dimensions in which language probability models can vary: (i) whether they make use of sequential information vs. hierarchical structure, and (ii) whether they predict word or parts-of-speech (POS). In [9], we showed that models of surprisal that only incorporate sequential information, whether of words or POS, fail to account for subtle distinctions in linguistic patterns.. The surprisal model that performed better in distinguishing the stimuli and replicating the expectation associated with the syntactic structure of a sentence was the one that considered hierarchical dependencies to predict the POS of the sentences, i.e., the syntactic surprisal. The hierarchical model that predicted individual words, i.e., the lexical surprisal, failed at replicating the same result. In [9], we concluded that surprisal models must therefore incorporate syntactic structure to mirror human listeners’ linguistic competence.

To evaluate whether syntactic surprisal modulations are similarly mirrored in brain data and provide an electrophysiological analysis of the theoretical conclusions reached in [9], in this paper, we used a new set of auditory stimuli containing homophonous sentences. In this new set of stimuli, the predictability of the syntactic content of the homophonous phrase is considered, allowing us to produce a high number of analytical contrasts among stimuli features (predictability, homophonous phrase type, and surprisal), to refine the knowledge of how syntactic information is processed in our brain, and how this neural processing is linked to the lexical and the syntactic surprisal.

We presented this new set of auditory stimuli to 32 healthy participants while recording their electroencephalographic (EEG) signal. We anticipate investigating into the correlation between various surprisal models and the syntactic modulation of our stimuli. We expect that each type of surprisal model has an influence on brain activity, even though in distinct manners and locations. We hypothesize that surprisal models incorporating both syntactic and morphological information will exhibit greater accuracy in discerning the neural activity associated with syntactic processing. This anticipation arises from our observation, supported by mathematical models, that these models uniquely discriminate our stimuli based on predictability [9]. Furthermore, we expect the temporal dynamics of the neural activity divergence between NPs and VPs to be highly affected by the predictability of the syntactic structure. Finally, we aim to replicate our prior findings using SEEG and a simplified set of stimuli, eliminating consideration for the predictability of syntactic structure [4], [6]. This comprehensive exploration promises to deepen our understanding of the electrophysiological correlates of syntactic processing.

## Methods

### Stimuli

To modulate the relation between the syntactic and surprisal information we crucially relied on the paradigm introduced in Greco et al. 2023 [9]. More specifically, three experimental conditions have been generated here by modulating the syntactic context preceding the HPs, which predicts the syntactic type of the HPs:

- **Unpredictable HPs** (Unpred.): the syntactic context preceding HPs is an adverb. Thus, the syntactic category of the HP is not predictable as the context allows both NPs and VPs. The syntactic category of the HP becomes discernible only after the HP: if it is followed by a verb, it is a NP (such as in *Forse* ***la porta*** *è aperta*, ‘Maybe **the door** is open’): otherwise, it is very likely a VP (*Forse* ***la porta*** *a casa*, ‘Maybe **s/he brings it** at home’). No differences will exist in the lexical surprisal values at the two HPs because the context preceding the HP is the same for both syntactic categories.
- **Strongly predictable HPs** (S. Pred.): the syntactic type of the HP is predictable at its onset. If the syntactic context preceding the HP is a verb the HP can only be a NP (such as in *Pulisce* ***la porta*** *con l’acqua*, ‘S/he cleans **the door** with water’). If the HP is preceded by a noun, it can only be a VP (*La donna* ***la porta*** *domani, ‘The woman* ***brings her*** *tomorrow’*). Our previous works [4], [6] exploited only this type of stimuli. The different lexical context preceding NPs and VPs allows for different lexical surprisal values.
- **Weakly predictable HPs** (W. Pred.): this is the mixed class. The sentences are introduced by a temporal adverb requiring a past tense (e.g., *Yesterday*). Thus, the first word of the HP (*la*) could either be an article (‘the’) or a clitic pronoun (‘her’), as in the Unpred. case; while the second word of the HP (*porta*) can only be a noun (‘door’), since the verbal form of the VP would involve a present tense verb ([s/he] ‘brings’) that is incompatible with the temporal adverb requiring a past tense (i.e. yesterday). An example stimulus is: *Ieri* ***la porta*** *era aperta*, ‘Yesterday **the door** was opened). Using this structure, in Italian, W. Pred. VP sentences are impossible, and thus W. Pred. HPs could only be NPs.

A total of 150 trials were prepared: 60 for Unpred. HPs, 30 NPs and 30 VPs, 60 for S. pred. HPs, 30 NPs and 30 VPs, and 30 for W. pred. HPs, only NPs since there cannot be VPs of this type.

### Surprisal calculation

Surprisal calculation is based upon on language probability models. Briefly, language probability models can be distinguished along two dimensions [9]:

- Structure: (i) Linear models (which we will instantiate as *n-gram models*) view language as an unstructured sequence. In such a model, the probability of an element determined by the linear sequence of *n* elements before it, and the probability of the entire sequence is the product of the individual elements. (ii) Hierarchical models (which we will instantiate as PCFGs), assume language is structured hierarchically, following Chomsky (1957) [11], so that probabilities are assigned to each element on the basis of the structural configuration it occurs, which can potentially span linearly unbounded distances.
- Prediction: (i) Word models predict individual words’ probabilities. (ii) Category models predict POS categories’ probabilities. Both can be easily computed using n-gram models. Roark et al. (2009) [10] show how word and category predictions can be separated in a hierarchical model, and we use their techniques to calculate syntactic surprisal for categories and lexical surprisal for words.

For subsequent analysis, we use only the surprisal values associated with the article (if NP) or clitic (if VP) of the HPs. As already demonstrated in [4, 6], this is the word for which the lexical surprisal difference between NPs and VPs is maximal for S. Pred. items.

### Human participants and EEG recordings

In total, 32 right-handed Italian native speakers were recruited (16 males and 16 females; median age 27, range 24-54). All participants retained the right to withdraw from the study at any time and received a small monetary compensation for their participation. They had no history of neurological or psychiatric conditions, normal hearing, normal or corrected vision, and no reported history of drug or alcohol abuse. All participants completed all experimental sessions. EEG recordings were carried out in a sound-isolation booth using a 65-channel HydroCel Geodesic Sensor Net (Electrical Geodesies, Inc., Eugene, OR). Electrode impedance was maintained below 30 kΩ throughout the recording session. Participants were instructed to listen carefully to the sentences to be able to answer questions about them. After reading instructions on a screen, they listened to the stimuli. The stimuli were administered four times, for a total of 600 trials. In each repetition block, the stimuli were presented in a randomized order. For each trial, the following events were annotated: the start of the sentence, the start of the HP, the start of the noun/verb of the HP, and the start of the first word following the HP. At the end of each repetition, participants were asked two questions about the stimuli (mean percentage of correct answers: 50%) (**Figure 1**). We do not consider accuracy in the task as important, as the task was presented to subjects to keep their attention high. The low accuracy is likely since questions were asked every 150 sentences. Participants were offered a break every 75 trials. The experiment lasted about one hour. EEG was acquired at 500 Hz. The present study received the approval of the Joint Ethics Committee of the Scuola Normale Superiore and the Scuola Superiore Sant’Anna (protocol n. 22/2022) and informed consent was obtained.

**Figure 1.**
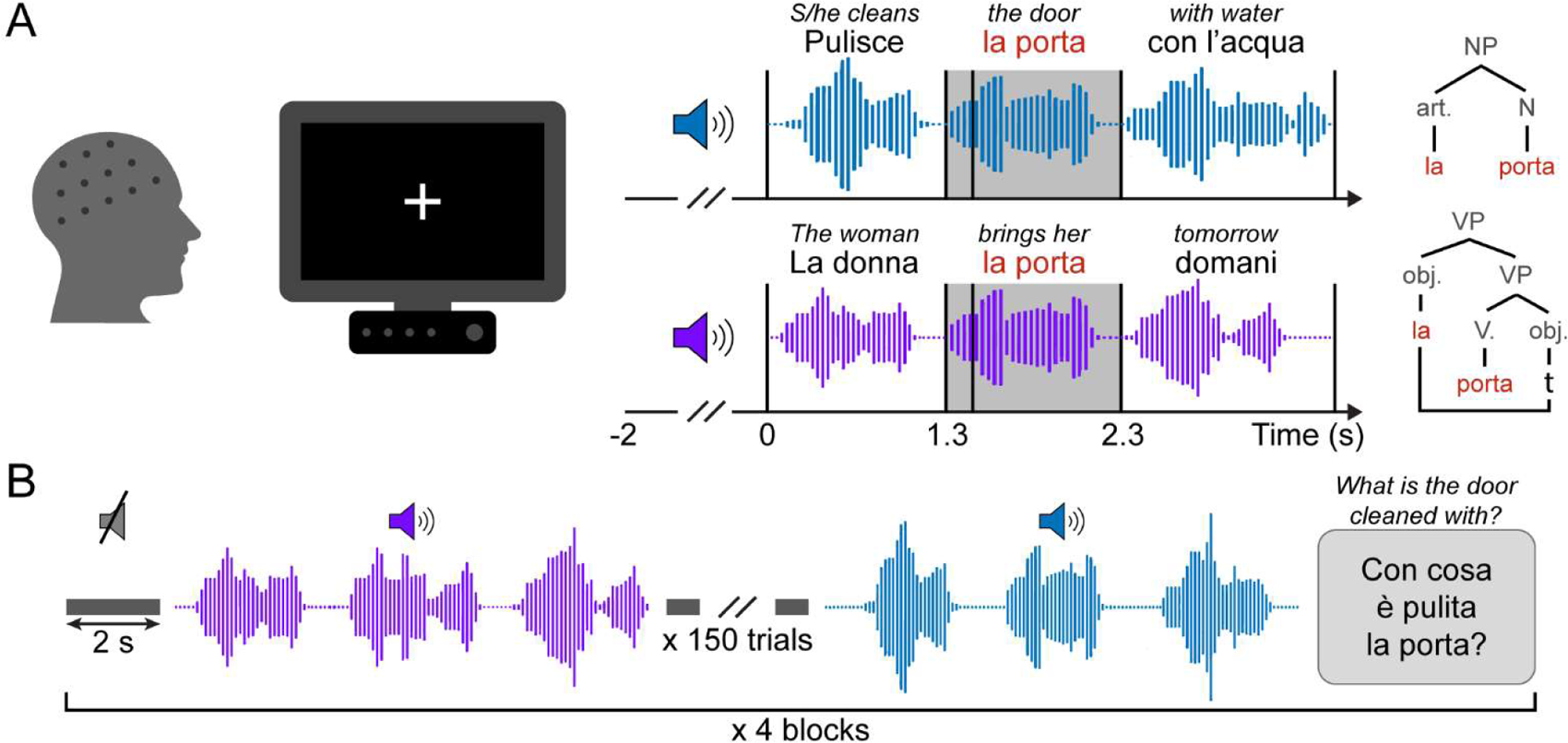
Recording protocol. **(A)** The EEG data were acquired while the participants fixated on a fixation cross displayed on a screen. After two seconds of silence, the sentence was played back through a speaker. For each sentence, its beginning, the beginning of the homophonous part, the beginning of the second word of the homophonous part, and the beginning of the word following the homophonous part were annotated. The black vertical lines on the graphs on the left represent the annotated events. The grey-shaded areas are the homophonous part. On top of each graph, there is an example stimulus sentence and its English translation. The top graph depicts an example of a strong predictable noun phrase, and the bottom graph shows a strong predictable verb phrase. The homophonous part (*la porta*) is highlighted in red. The syntactic trees of the example stimuli are drawn on the right of the graphs. *t* indicates the position where the pronoun is base generated in the verb phrase. **(B)** Data acquisition was divided into 4 blocks. In each block, the 150 stimulus sentences were presented in a randomized order, with 2 s of silence between one stimulus and the other. At the end of each block, the participants were asked to answer two questions regarding the content of the stimuli.

### EEG pre-processing

First, EEG data were downsampled at 250 Hz to reduce computational time. Then, 2 pre-processing steps were carried out, using a semiautomatic pipeline [12], [13], [14]. The pre-processing was divided into two steps to minimize the removal of brain activity while maximizing the quality of the Independent Component Analysis (ICA) decomposition.

#### Pre-processing step 1

EEG data were band-pass filtered at (1 – 40 Hz) using a Hamming windowed sinc FIR filter [15]. Then, the signal was divided into non-overlapping windows of length equal to 1 s. The windows with a joint probability larger than three standard deviations with respect to the mean probability of occurrence of a trial were rejected. Independent Components (ICs) were calculated using the Infomax algorithm [16]. Finally, ICs representing notable eye artifacts were rejected [17].

#### Pre-processing step 2

Final ICA weights resulting from Pre-processing Step 1 were applied to data more conservatively pre-processed within EEG Pre-processing Step 2. The data were epoched from 2 s before the stimulus onset to 4.5 s after and band-pass filtered (0.1 – 40 Hz) using a Hamming windowed sinc FIR filter [15]. Epoch length was chosen to always contain the entire stimulus. Bad channels and epochs containing high-amplitude artifacts, high-frequency noise, and other irregular artifacts were removed. Finally, bad channels were interpolated using cubic-spline interpolation [18] and the EEG data were re-referenced to the average.

### Event-related spectral perturbation estimation

Previous work showed that the frequency of the EEG signal activity plays an important role in syntactic processing [4], [6], [19], [20], [21]. Thus, we computed event-related spectral perturbations (ERSPs) to characterize the neural response to our stimuli in both frequency and time. Time-frequency transforms of each trial were normalized to the baseline (divisive baseline, ranging from -2000 ms to 0 ms before the start of the sentence), time-warped to the stimulus events [4], [22], and averaged across trials for each participant, for each stimulus class to obtain the ERSPs [23].

### Representational similarity analysis

Representational similarity analysis (RSA) is an analysis technique used to compare the information content carried by a representation in the brain with that carried by a model [24]. This is done by comparing the representational dissimilarity matrices (RDMs) of brain activity with those computed on some features of the stimuli. The RDMs are square matrices of pairwise dissimilarity values for all pairs of stimulus-specific patterns.

More specifically, the steps of the RSA analysis are:

- ***Computation of model RDMs.*** The model RDMs are calculated on some features of the stimuli and not on the EEG data. They were calculated on several dimensions of our stimuli: the phrase type (NP or VP), the predictability (S. pred., W. pred., Unpred.), the lexical surprisal of the article/clitic of the HP, the syntactic surprisal of the article/clitic of the HP, the n-gram surprisal of the article/clitic of the HP, and the POS n-gram surprisal of the article/clitic of the HP (**Figure 2A** and **Figure 4A**). For the surprisal values, the RDMs were calculated using the Euclidean distance between the pairs of averages of the surprisal values for a given stimulus class. Having a total of 5 classes (Unpred. VPs and NPs, S. Pred. NPs and VPs, and W. Pred. NPs), the RDMs are 5x5 matrices. The value in row *i* and column *j* of the RDMs for the lexical and the syntactic surprisal is the difference between the mean value of the lexical (or syntactic) surprisal across all the stimuli of class *i* and the mean value of the lexical (or syntactic) surprisal calculated across all the stimuli of class *j*. The lexical surprisal and syntactic surprisal RDMs are thus matrices composed of continuous real values. For the phrase type RDM, the Hamming distance was used, resulting in a binary RDM. The value in row *i* and column *j* of the RDM for the phrase type is 0 if the phrase type of class *i* is the same as the phrase type of class *j*, 1 otherwise. The predictability RDM is a 3-valued matrix, with the distance between items belonging to the same class being 0, the distance between S. pred. and Unpred. is 1, and the distance between W. pred. and the other two classes is 0.5. The distance value of 0.5 was chosen because W. pred. is an intermediate class between S. pred. and Unpred.
- ***Computation of brain* RDMs.** To calculate brain RDMs, the 5 condition-specific ERSPs were windowed in the time domain with a window length of 200 ms and an overlap of one time sample (4 ms at 250 Hz). The pairwise Euclidean distance between the ERSPs time-frequency samples of two analogous time windows was then calculated for each pair of conditions. This procedure was repeated for each time window, for each frequency, for each channel, and for each participant, resulting in channel-time-frequency-varying participant-specific RDMs.
- ***Comparison between brain RDMs and model RDMs.*** The brain RDMs were then compared to the model RDMs using the correlation coefficient as an index of similarity, resulting in a correlation value for each time-frequency point, for each model, for each channel, and for each participant.
- ***Statistical analysis on the correlation values.*** Finally, these correlation values were tested against the null hypothesis of being equal to 0 (see Statistical analysis).

### Linear modeling

For each participant, we modeled the ERSPs using linear regression. The linear regression aims to model the observed neural response (time-frequency point, for each channel, for each participant) as a linear combination of different features of the stimuli. The features of the stimuli (model regressors) used were: the phrase type (NP or VP), the predictability (S. pred., W. pred., or Unpred.), their interaction (phrase type : predictability), the lexical surprisal of the article/clitic of the HP, and the syntactic surprisal of the article/clitic of the HP. QR decomposition was used to solve the linear model and find the regression coefficient for each regressor [25]. Specifically, the time-warped trial-by-trial time-frequency transforms of the EEG signal were used to estimate the regression coefficients for each participant, for each time-frequency point, and for each channel, resulting in 4 (one for each regressor) 4-dimensional regression coefficients. Finally, these regression coefficients were tested against the null hypothesis of being equal to 0 (see Statistical analysis). We repeated the analysis by deleting the syntactic surprisal regression term to avoid redundancy between this and the interaction between the phrase type and predictability. These two terms, by design, should convey the same information [9].

### Statistical analysis

Both the correlation coefficients of the RSA, and the regression coefficient of the linear modeling have 4 dimensions: time, frequency, channels, and participants. Thus, it is possible to perform a cluster-based permutation test in the time-frequency-spatial domain [26].

The participant-specific correlation values of RSA were compared against matrices of zeros of the same size to test for the null hypothesis of zero correlation.

To increase the power of the statistical test, the regression coefficients of the linear model were first averaged in 5 frequency bands (delta: 0-4 Hz, theta: 4-8 Hz, alpha: 8-13 Hz, beta: 13-30 Hz, and gamma: 30 – 40 Hz) and 3 time windows (from the start of the sentence to the start of the HP, the HP, and from the end of the HP to the end of the sentence). The windowed regression coefficients were thus compared against the null hypothesis of their value being equal to 0, similar to the correlation values of the RSA [27].

Pair-wise comparisons of NPs and VPs trials were carried out in the same way, but directly on the time-frequency transforms of the EEG data. Pair-wise comparison of S. pred. and Unpred. items were computed using a cluster-based permutation test directly on the time-warped pre-processed EEG signal.

## Results

### Syntactic surprisal and syntactic class are represented by neural activity

Representational similarity analysis (RSA) was performed to investigate the effect of four dimensions of our stimuli [the lexical surprisal, the syntactic surprisal, the phrase type (NP or VP), and the predictability (S. Pred., W. Pred, and Unpred.)] on the brain response, as coded by the ERSPs (i.e. time-frequency transforms).

First, we found that ERSPs power did not correlate with lexical surprisal. One cluster of significant negative correlation was found for the syntactic surprisal in the gamma band during the presentation of the homophonous parts of the stimuli. This negative correlation was found between frontal-left electrodes (**Figure 2B**, top row).

For the phrase type, one cluster of significative negative correlation between the brain RDM and the model RDM was found. Right electrodes responded to the phrase type just after the start of the sentences, in theta band. The significance lasted up until the start of the second word of the homophonous part of the stimuli (noun or verb) (**Figure 2B**, second row).

For the predictability, two clusters of significant negative correlation were found: (i) frontal electrodes (with no evident lateralization) responded to the predictability of the stimuli after the start of the second word of the homophonous part (noun or verb), in the alpha band; (ii) frontal electrodes (with slight lateralization to the right) significantly correlated with the predictability RDM after the start of the first word following the homophonous part, with the significance being in a frequency band between alpha and beta. (**Figure 2B**, graphs three and four, from the top, one for each significant cluster).

**Figure 2.**
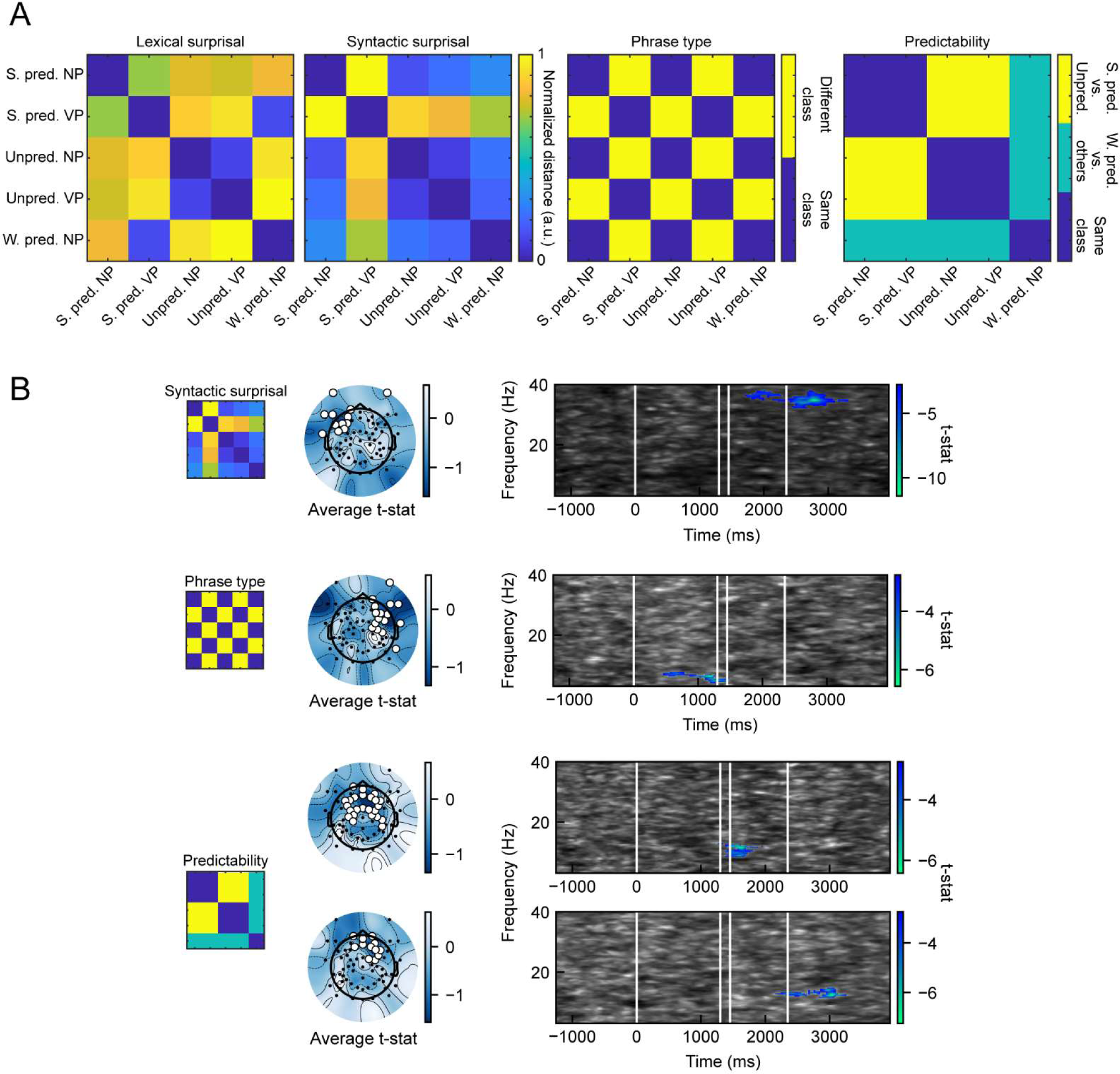
Representational similarity analysis. (A) Models used in representational similarity analysis. Each of the four matrices is called Representational Dissimilarity Matrix (RDM). Each RDM is a different representation of our stimuli, along 4 dimensions: the hierarchical lexical surprisal, the syntactic surprisal, the phrase type (NP or VP), and the predictability (S. pred, W. pred, or Unpred.). (B) Significant clusters for the representational similarity analysis. The topographic plots show the average t-statistic across significant time points and frequencies. White dots represent the significant electrodes. The time-frequency graphs represent the minimum t-value across significant electrodes. Significant time-frequency points are colored in blue (top graph for the syntactic surprisal, second graph for the phrase type, and graphs three and four for the two significant clusters for the predictability). The time is adjusted according to the stimulus onset (0 ms). The four white vertical lines respectively represent: (i) stimulus onset, (ii) the start of the article/clitic, (iii) the start of the noun/verb, (iv) the start of the word that follows the noun/verb.

### The neural response to the syntactic class is not dependent on lexical surprisal

RSA is not a multivariate analysis, i.e., treating all the models independently, it is not able to decouple the effect of confounding factors such as the lexical surprisal from the neural response to features of interest, such as the phrase type. Thus, we performed linear modelling, a multivariate analysis, on the ERSP power values with the syntactic surprisal, lexical surprisal, phrase type, predictability, and the interaction between phrase type and predictability as regressors. This analysis revealed only an effect of the lexical surprisal. A new linear model was treated without the syntactic surprisal under the hypothesis that having both the syntactic surprisal and the interaction between predictability and phrase type in the linear model is redundant [9]. This redundancy may cause to decrease the single effects of the syntactic surprisal and the interaction term making them (singularly) not significant. We chose to utilize the interaction between phrase type and predictability instead of syntactic surprisal because the term ’interaction’ precisely describes our stimuli. Investigating the interplay between phrase type and predictability allows us to discern syntactic computation in the brain from other types of language processing. **Figure 3A** shows the results of the linear modeling without the syntactic surprisal term. This analysis revealed: (i) a significant negative regression coefficient for the phrase type in the beta band, during the homophonous part of the sentences, in central and left electrodes; (ii) a significant positive regression coefficient for the interaction between phrase type and predictability in the beta band, during the homophonous phrases, in central and frontal electrodes; and (iii) a significant negative regression coefficient for the lexical surprisal in the delta band, during the start of the sentences and their homophonous parts, in posterior electrodes. Importantly, lexical surprisal correlates with brain activity in a different frequency band, a different time window, and different electrodes than the interaction term.

### Syntactic predictability modulates the response to NPs and VPs

Figure 3B displays the results of the cluster-based permutation test on ERSPs for the interaction between phrase type and predictability, broken down by predictability. For S. pred. sentences, the contrast between NPs and VPs revealed a stronger negative response for VPs, in the beta band (beta desynchronization), during the homophonous part of the stimuli. This stronger beta desynchronization for VPs was found in the central and right electrodes. For Unpred. sentences, the difference between VPs and NPs was found in lower frequency bands, after the homophonous phrases, where syntactic interpretation is hypothesized to be carried out by the participants. The response to VPs was stronger than the response elicited by NPs, in the delta band, for almost all the EEG recording contacts. In theta band, response to Unpred. VPs was stronger than the response to Unpred. NPs on the channels over the left hemisphere.

**Figure 3.**
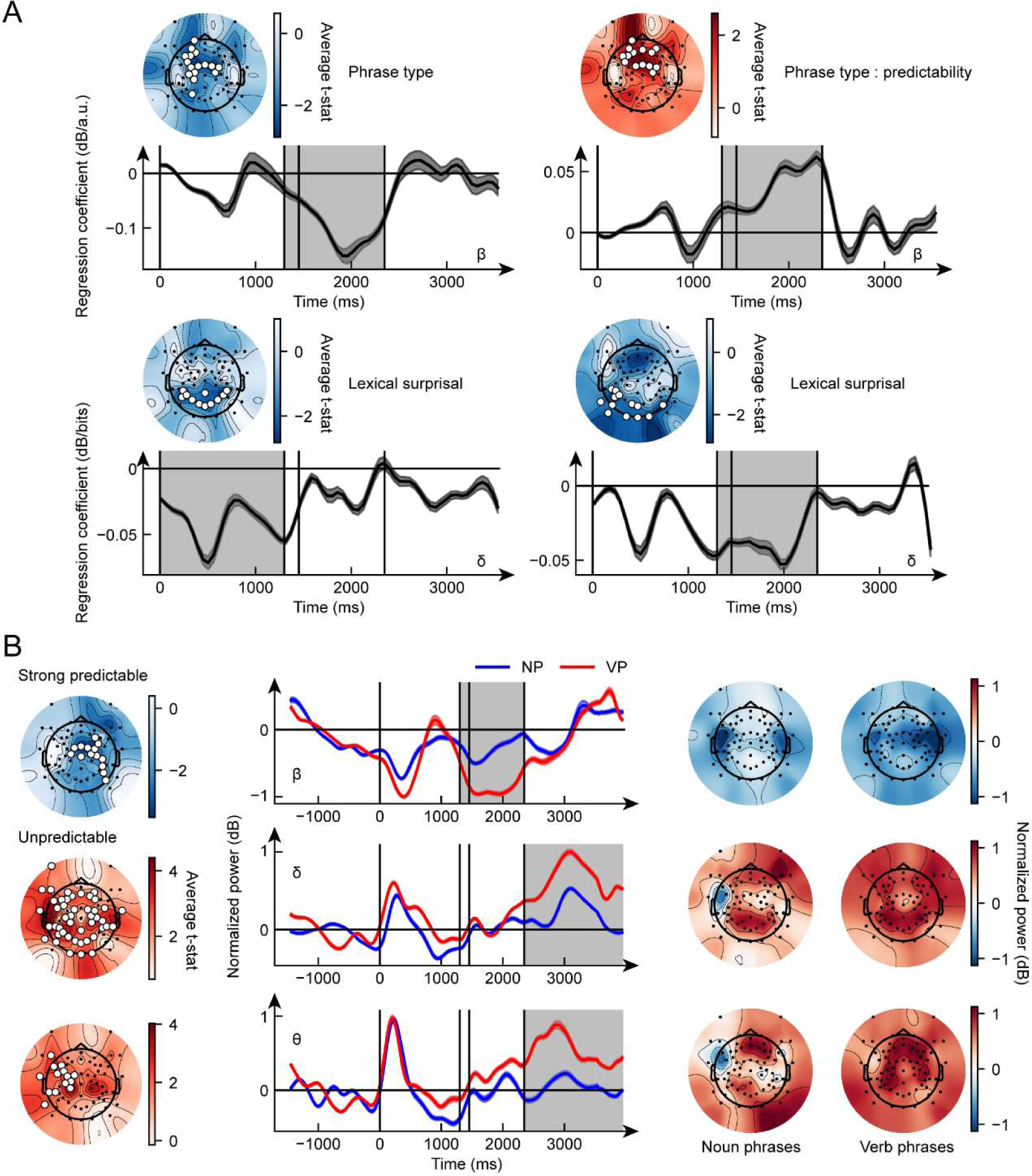
Noun phrases vs. verb phrases. (A) Significant clusters (p < 0.05) for the linear modeling analysis. The topographic plots (head plots) show the average t-statistic across significant time points and frequencies. White dots represent the significant electrodes. The line graphs represent the temporal evolution of the coefficients of the linear model, averaged across significant electrodes, for the given frequency band. The dark grey shaded area represents the standard error across participants, light grey shaded area shows the significant time points. The time is adjusted according to the stimulus onset (0 ms). The four black vertical lines respectively represent: (i) stimulus onset, (ii) the start of the article/clitic, (iii) the start of the noun/verb, (iv) the start of the word that follows the noun/verb. The interaction between phrase type and predictability is denoted as phrase type : predictability. (B) Results of the cluster-based permutation test on the ERSPs for the contrast noun phrases vs. verb phrases, for the strong predictable items (top row) and the unpredictable items (second and third rows). No contrast was done on weakly predictable items since there were only noun phrases. Topographic maps in the first column are the same as in (A). The line graphs represent the temporal evolution of the power of the ERPSs averaged across significant electrodes and for the given frequency bands. The last two columns represent the average power across significant time points and participants, for the given frequency band, for the sentences containing noun phrases (left) and verb phrases (right).

### Linear surprisal models do not affect syntactic neural computation

We repeated the RSA and the linear modelling analysis account also for the linear models of surprisal, i.e., the n-gram surprisal, and the POS n-gram surprisal (Figure 4A). The RSA revealed that the POS n-gram surprisal (unlike the syntactic surprisal) is not represented by the electrophysiological activity. The n-gram surprisal showed one cluster of significant negative correlation in the gamma band during the presentation of the homophonous parts of the stimuli and at the end of the sentences. This negative correlation was found between frontal-left electrodes (Figure 4B) and only partially overlaps with that associated to the syntactic surprisal (Figure 2B).

The n-gram surprisal values and the POS n-gram surprisal values were added to the regressor for the linear modeling. Thus, the regressor for the final linear model were: n-gram surprisal, POS n-gram surprisal, lexical surprisal, phrase type, predictability, and the interaction between phrase type and predictability. Both n-gram surprisal and POS n-gram surprisal significantly drive EEG activity. The POS n-gram surprisal has an effect at the start of the sentence (before the homophonous part) in delta and beta bands (Figure 5, top 2 graphs). The n-gram surprisal has an effect at the start of the sentence (before the homophonous part) in delta band (Figure 5, bottom graph).

**Figure 4.**
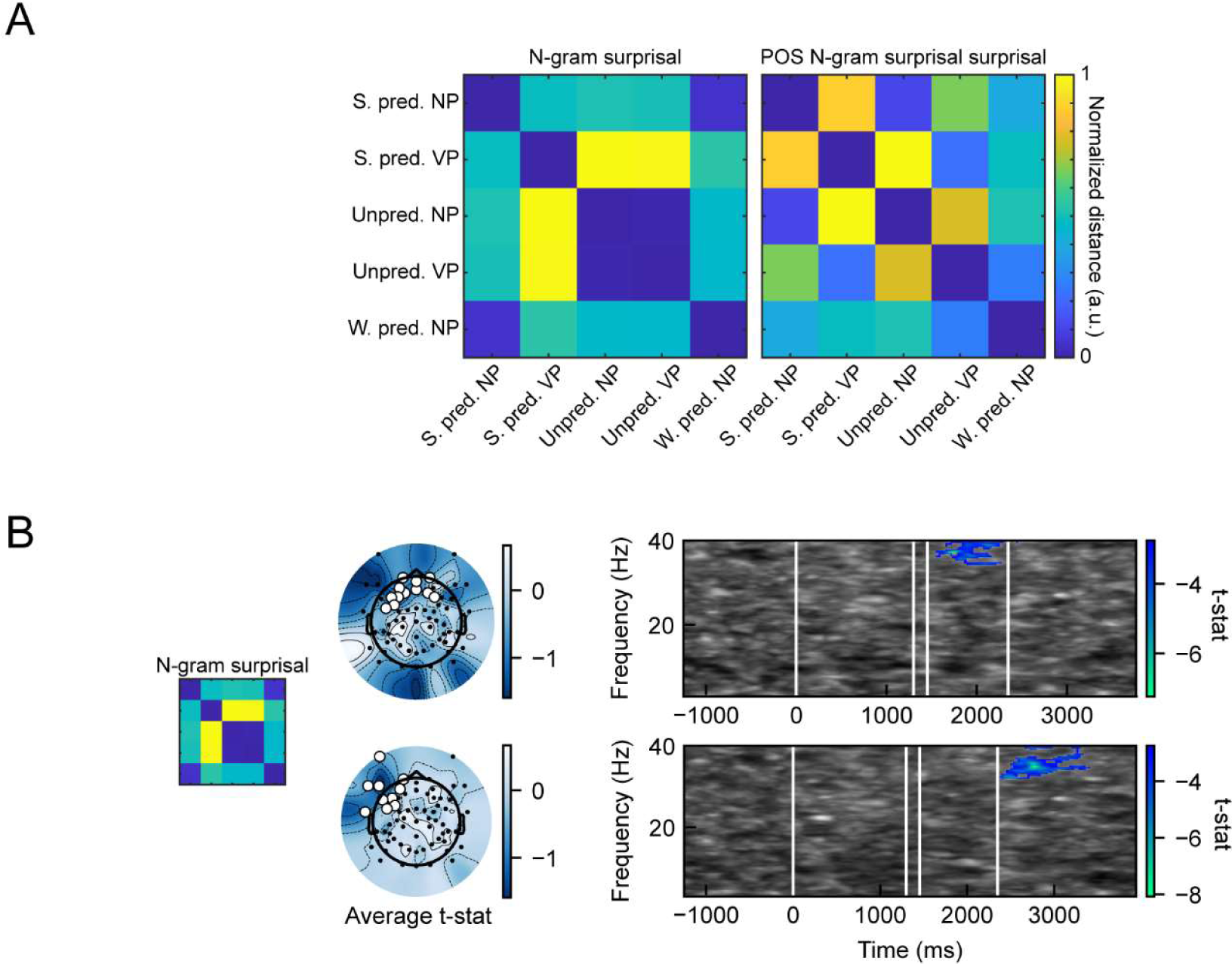
Representational similarity analysis of the linear surprisal. (A) Models used in representational similarity analysis. Each of the two matrices is called Representational Dissimilarity Matrix (RDM). Each RDM is a different representation of our stimuli, along two dimensions: the n-gram surprisal, and the POS n-gram surprisal. (B) Significant clusters for the representational similarity analysis. The topographic plots show the average t-statistic across significant time points and frequencies. White dots represent the significant electrodes. The time-frequency graphs represent the minimum t-value across significant electrodes. Significant time-frequency points are colored in blue (both graph for the n-gram surprisal, for the two significant clusters). The time is adjusted according to the stimulus onset (0 ms). The four white vertical lines respectively represent: (i) stimulus onset, (ii) the start of the article/clitic, (iii) the start of the noun/verb, (iv) the start of the word that follows the noun/verb.

**Figure 5.**
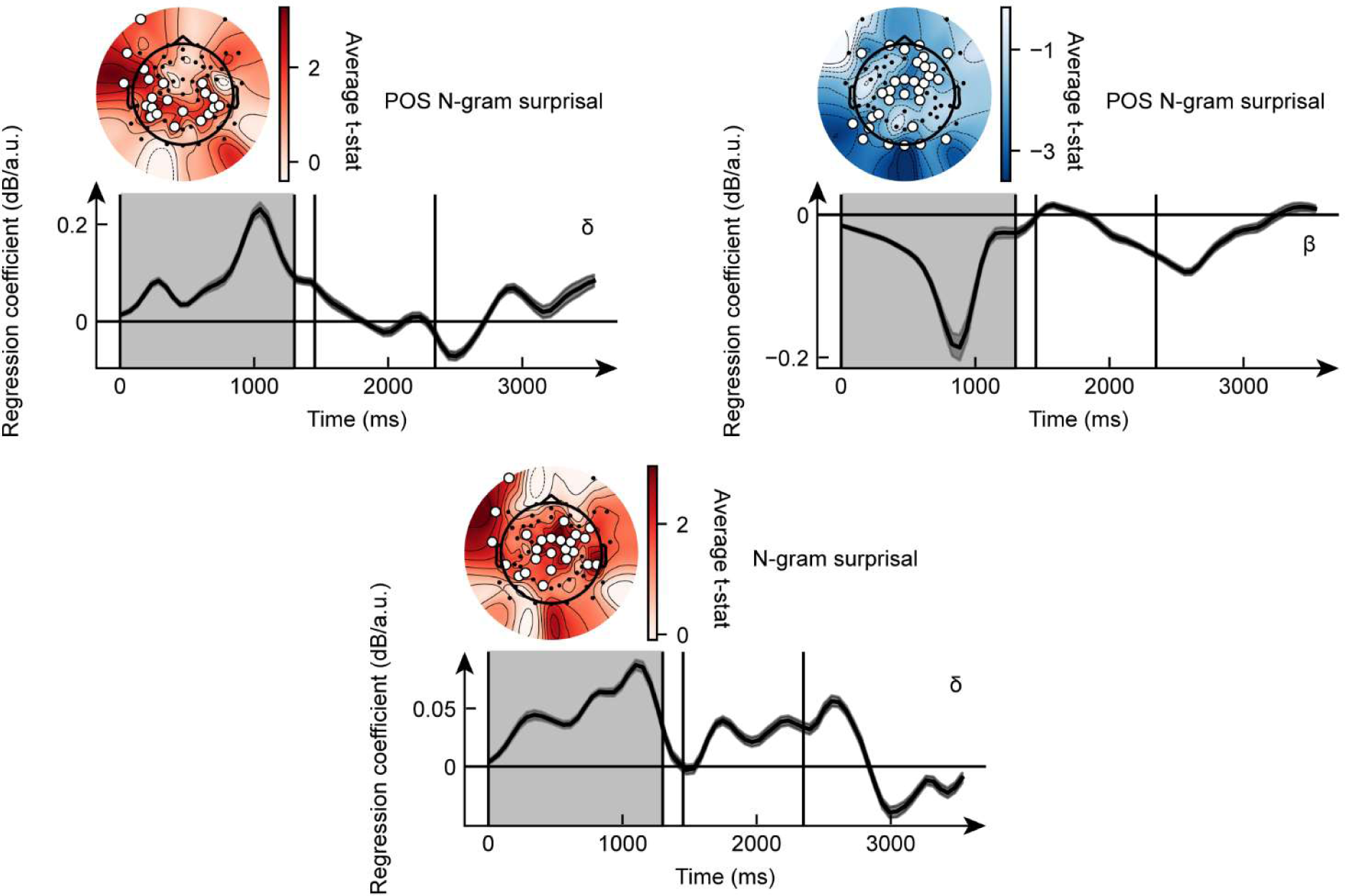
Linear modeling analysis with linear surprisal. Significant clusters (p < 0.05) for the linear modeling analysis. The topographic plots (head plots) show the average t-statistic across significant time points and frequencies. White dots represent the significant electrodes. The line graphs represent the temporal evolution of the coefficients of the linear model, averaged across significant electrodes, for the given frequency band (Greek letter in the graph). The dark grey shaded area represents the standard error across participants, light grey shaded area shows the significant time points. The time is adjusted according to the stimulus onset (0 ms). The four black vertical lines respectively represent: (i) stimulus onset, (ii) the start of the article/clitic, (iii) the start of the noun/verb, (iv) the start of the word that follows the noun/verb.

### Neural response to syntactic category does not depend on systematic confounding effects

To exclude systematic differences in potential lexico-semantic confounding factors between NPs and VPs, we used a cluster-based permutation test directly on the pre-processed EEG signal and defined two contrasts: S. pred. NP vs. Unpred. NP, and S. pred. VP vs. Unpred. VP. The rationale behind this analysis is that if we find the same significant effects for both contrasts, then the systematic differences between S. pred. NPs and S. pred. VPs that could have affected the previous results can be excluded. We do not expect systematic differences in confounding factors in Unpred. NPs and Unpred. VPs since the structure of Unpred. sentences do not allow them by design. We performed this analysis directly on the filtered data, and not on the time-frequency transformed EEG. This choice allowed us to explore the differences occurring in shorter time windows than those possible with ERSPs, due to the dilution effect of the filtering on the signal. Indeed, we expect those systematic differences to occur during short time windows.

For the S. pred. NP versus Unpred. NP contrast, 10 clusters were found. For the S. pred. VP vs. Unpred. VP 14 clusters were found. No differences were found during the baseline period. Six pairs of clusters were comparable across the two contrasts (Figure 6). Among these six pairs, two spatio-temporal significant clusters were found during the start of the sentence: one involving frontal, temporal, and posterior electrodes, denoting a higher positive potential peak for Unpred. sentences right at the start of the stimuli (Figure 6, first row); the other involving central electrodes and denoting a negative potential deflection for Unpred. sentences, absent in S. pred. stimuli, right before the start of the homophonous phrase (Figure 6, second row). Two significant clusters, with opposite polarity, were found in the frontal and posterior regions during the homophonous phrases, indicating a stronger potential deflection for S. pred. sentences (Figure 6, third and fourth rows). The last two significant clusters, again with opposite polarity, were found during the last part of the sentences, in frontal and posterior electrodes, respectively (Figure 6, fifth and sixth rows).

**Figure 6.**
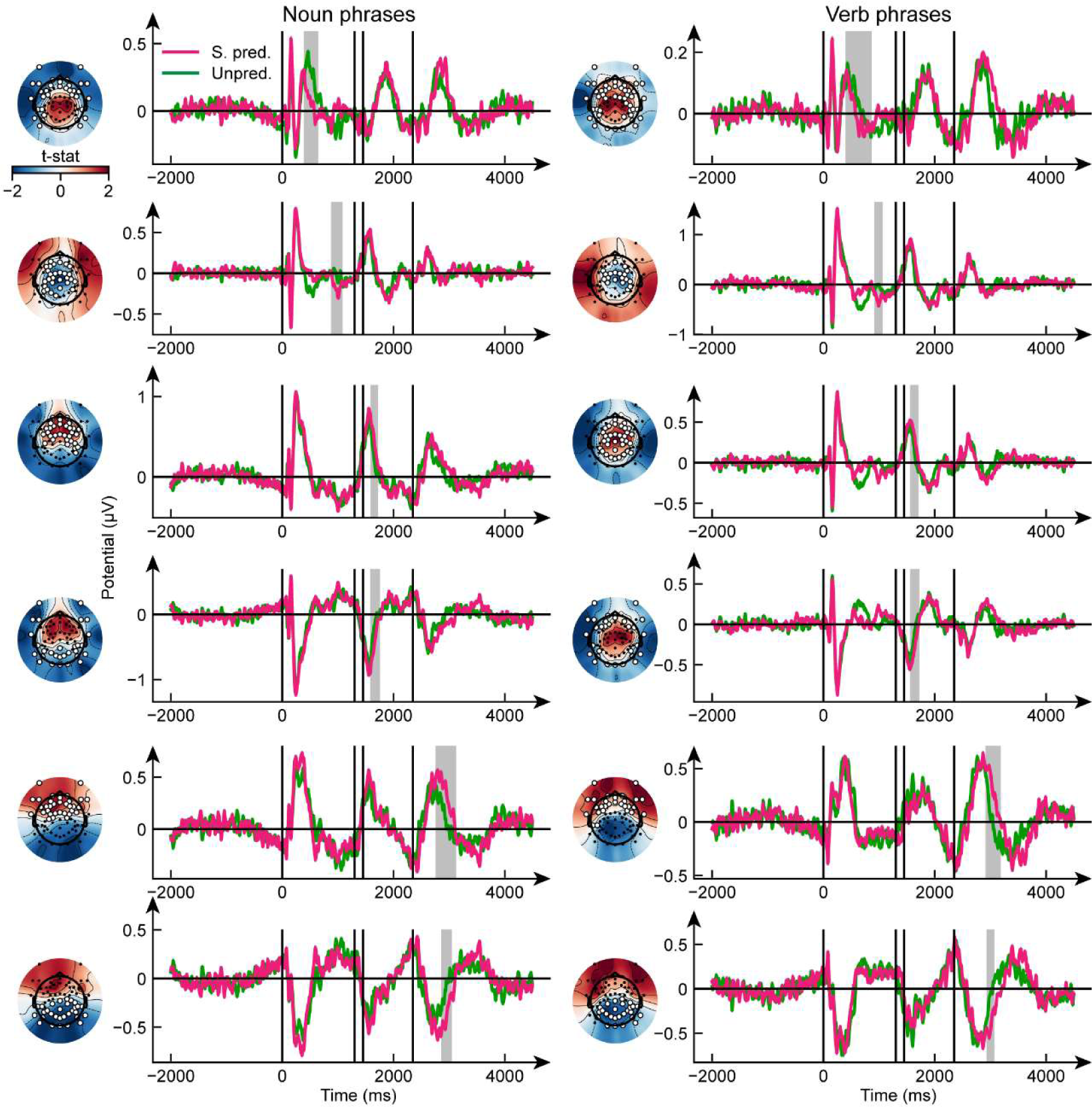
Strongly predictable vs. Unpredictable sentences Results of the cluster-based permutation test on ERPs for the contrasts S. pred. NP vs. Unpred. NP (left) and S. pred. VP vs. Unpred. VP (right). The topographic plots show the average t-statistic across significant time points. The line graphs represent the temporal evolution of the EEG signal, averaged across participants and significant electrodes. The light grey shaded area shows the significant time points (cluster-based permutation test). The time is adjusted according to the stimulus onset (0 ms). The four black vertical lines respectively represent: (i) stimulus onset, (ii) the start of the article/clitic, (iii) the start of the noun/verb, (iv) the start of the word that follows the noun/verb.

## Discussion

In our earlier research [4], [6], SEEG recordings were used to compare the brain activity elicited by the processing of (homophonous) noun phrases or verb phrases. The primary neural correlate of syntactic processing was found in the increase in power of frequencies in the high-gamma band (150-300 Hz). We discovered that there were more responsive contacts for VPs than for NPs, with the neural network supporting VP processing being wider and involving more cortical and subcortical areas than the network processing NPs. Unfortunately, the concept of n-gram surprisal represented a confounding factor for these findings. The higher the rarity of a sequence of words, the higher the surprisal, which is defined as the negative log probability of a given word following another in a sentence given a corpus [7]. It is well known that surprisal and brain activity are positively correlated [8], and VPs were associated to a generally higher n-gram surprisal than NPs. Here, we modulated both the syntactic and lexical surprisal values of the HPs, showing that the difference in the neural processing of NPs or VPs is better mirrored by the syntactic surprisal.

### Lexical predictability does not affect syntactic processing

Here we used two models of surprisal calculated using hierarchical structures: the lexical surprisal (word prediction), and the syntactic surprisal (hierarchical POS prediction). Given the nature of the two surprisal measures and the fact that only the syntactic surprisal was able to fully characterize our stimuli [9], our results are driven by the syntactic predictability of the HPs rather than their lexical predictability. On the other hand, the neural correlates of lexical surprisal may be related to other aspects of language processing, such as processing difficulty during language comprehension [28], [29], and semantic information retrieval [30]. Moreover, the RSA showed that lexical surprisal failed at eliciting a significant brain response thus that our stimuli were able to induce syntactic computation while controlling for other types of language computation. The linear model showed an effect of the lexical surprisal in posterior electrodes, in the delta band, from the start of the sentence to the end of the homophonous part. This result aligns with the large body of evidence demonstrating a crucial role of temporo-parietal areas for language comprehension [31], [32], [33]. Moreover, the prominence of delta oscillations in this context could be attributed to their recognized role in orchestrating long-range communication and coordinating cognitive processes, such as memory retrieval and attention allocation [34], [35]. This convergence of findings substantiates the notion that lexical surprisal exerts a distinct and lasting impact on neural processing. Importantly, this impact does not overlap with the neural activity related to syntactic processing, as the latter is prominent in different electrodes, different frequency bands, and different time windows.

### The electrophysiological correlates of NPs and VPs

Phrase type was represented by the observed differences in neural patterns during the start of the sentences, and the homophonous part. The significant correlation at the start of the sentence may be due to the presence of verbs and nouns before the homophonous part for the S. pred. NP and S. pred. VP stimuli, respectively. Verbs and nouns in isolation are known to elicit different brain responses [36], [37], [38]. However, we introduced the concept of predictability for this purpose: eliminating confounding factors thanks to the number of contrasts that we can obtain by comparing our stimuli along different dimensions.

To this aim, we used the linear modeling analysis to isolate the effects of the phrase type, the predictability, their interaction, and the lexical surprisal. We found an effect of the phrase type in the beta band, during the homophonous part, on central/left electrodes. The effect of the interaction between the phrase type and the predictability was in central/frontal electrodes in the beta band during the homophonous part. We broke down the effect of the phrase type by the predictability and we found that: (i) for the S. pred. items, VP elicited higher beta desynchronization during the homophonous part, over central/right electrodes; and (ii) in the Unpred. case, VPs caused a delta (over most electrodes) and theta (on right electrodes) synchronization after the homophonous part of the sentence. The lack of a significant interaction effect between the phrase type and the predictability modeled with linear regression may be attributed to the lower statistical power in comparison with this post-hoc comparison. The fact that in the S. pred. case VPs elicited higher brain activity than NPs, during the homophonous part confirmed our previous findings using SEEG [4]. In this earlier study, only S. pred. stimuli were used. Furthermore, SEEG and EEG signals have different characteristics. For instance, the high-gamma band is not recordable using EEG due to the low-pass filtering effect of the scalp and the skull [39], and future work should further establish whether there is a direct link between the beta band desynchronization in EEG and high-gamma increase in SEEG. In summary, although expanding on our previous work, we have also confirmed with EEG data what we previously observed using SEEG recordings, opening new research and treatment possibilities as EEG is much simpler to record than SEEG, and comes with fewer limitations. However, if spatial resolution is of utter importance for the investigation of the neural correlates of syntactic processing, SEEG still has an advantage [42], and future work is needed to precisely identify the cortical areas whose activity correlates with the syntactic predictability.

The latency of the difference between NPs and VPs in the Unpred. scenario (i.e., after the homophonous part) is consistent with the hypothesis on the timing in which the participant realizes what type of phrase has been heard (NP or VP). Thus, the detected activity in delta and theta bands may be an index of late disambiguation. It is not surprising that we found different bands for the two predictability scenarios. In the case of S. pred. sentences, evidence of disambiguation is absent. This is because participants know if they are listening to a NP or a VP prior to the onset of the homophonous part. On the other hand, in the Unpred. case, participants must remain uncertain about the structure of the sentence following the homophonous phrase and should continue to show evidence for disambiguation. Thus, the late response in delta-theta bands probably reflects this process. The shift of electrophysiological activity in lower frequencies, together with the higher number of selective electrodes, may be a correlate of the higher syntactic processing effort required from the listener in the Unpred. scenario [34], [40], [41].

Of note, the unpredictable sentences shift the significant activity in the portion that follows the homophonous phrase, thus the attribution of the class of the HP (NP or VP) takes place after the phrase has been listened to.

The significant clusters for the contrasts S. pred. NP vs. Unpred. NP, and S. pred. VP vs. Unpred. VP are highly superimposable. This shows that there are no systematic differences in potential confounding factors between NP and VP stimuli, in both the S. pred. sentences and in the Unpred. sentences. Thus, the results shown in Figure 3B and section **Syntactic predictability modulates the response to NPs and** VPs are only due to the effect of the different neural activities underlying the processing of different syntactic structures.

We did not perform any direct contrast involving W. pred. sentences because W. pred. VP sentences are impossible in Italian and thus the responses to NPs vs. VPs could not be compared. However, their inclusion in the experiment was fundamental for the finer modulation of syntactic surprisal values that we achieved. Without W. pred. sentences we would have less variety in the values of surprisal and therefore a more limited ability to infer what is happening in the brain due to the values of surprisal. Nevertheless, W. pred. sentences open many possibilities for future studies.

### Syntactic processing vs. surprisal

Syntactic surprisal has a significant effect on the neural response, especially in beta and gamma bands. The electrophysiological response to the syntactic class depends on the predictability of the syntactic class, and thus on syntactic surprisal, but not on lexical surprisal, indicating that the observed response is necessarily syntactic.

Our previous paper [9] has established that the predictability associated with three distinct classes of stimuli is more accurately represented in a model that incorporates syntactic information. This assertion is further corroborated by our recent findings. However, the RSA analysis conducted on the linear surprisal values revealed no significant role of surprisal models based solely on POS. This suggests that the morphological information encoded in POS alone may be insufficient, and syntax is indeed necessary to elucidate brain activation patterns. If validated, this would serve as additional evidence supporting our hypothesis.

Conversely, our analysis identified a significant role of surprisal based on n-gram models. This aligns with expectations, given the numerous studies demonstrating the impact of n-gram-based surprisal on cognitive tasks. However, our previous paper on surprisal indicated that n-gram models are not optimal for distinguishing between different classes of stimuli in terms of predictability. Therefore, it is not concerning that we observed activations correlating with both n-gram surprisal models and syntactic models. Importantly, the electrodes that were significant in the two conditions only partially overlapped, suggesting that they represent different facets of linguistic stimuli processing.

These results confirm the pivotal role of the computation of syntactic structures in human languages [42]. We showed different roles for the various EEG frequency bands [43], showing an immediate response that is syntactic specific in the beta band in the contrast between S. pred. sentences and a late response in delta and theta bands in the contrast between unpredictable items. This delta-theta response is an index of disambiguation of the syntactic type of the homophonous phrase after that the phrase type becomes discernible, coherently with the higher cognitive load associated with syntactic processing for the Unpred. items. Overall, our findings suggest that the processing of noun phrases and verb phrases is modulated by the syntactic surprisal as encoded by the predictability of the HPs and that there are distinct neural representations for strongly syntactically predictable and syntactically unpredictable stimuli (Figure 7).

**Figure 7.**
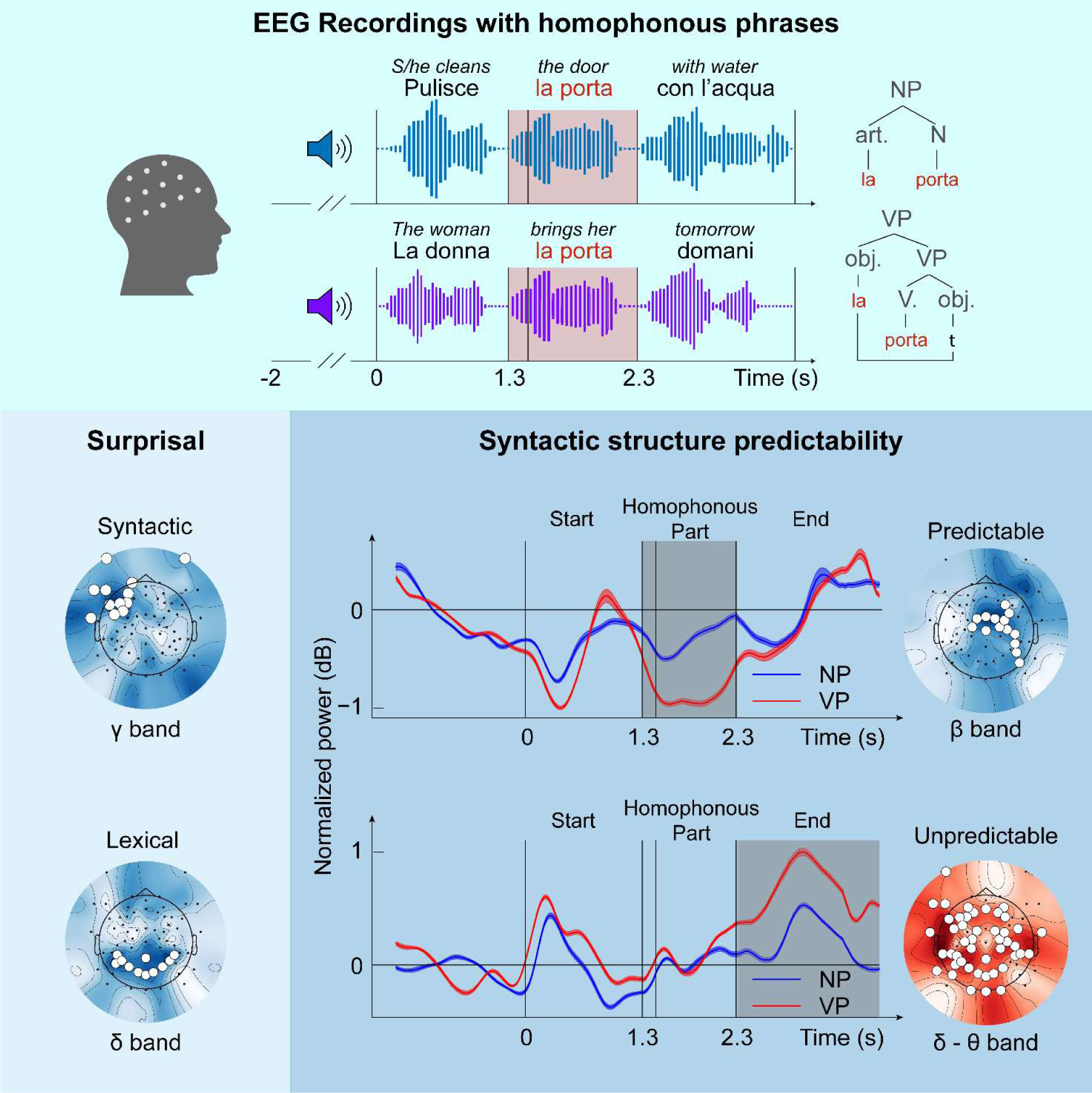
Main Results. Graphical summary of the results presented in this paper. Refer to the other figures for the panel legends.

## Conclusions

This paper showed that the class of predictability correlates with brain activity as predicted by mathematical models [9]. The observed responses are inherently syntactic because of the distinctions in NPs and VPs contrasts in S. pred. and Unpred. cases, the alignment of their temporal dynamics with the expected syntactic processing timings, and the exclusion of acoustic factors, coupled with the integration of surprisal and syntactic processing concepts.

In this sense, these results constitute a fundamental factor of a broader picture toward the cracking of the syntactic code of human languages. More specifically, the present results strongly correlate and provide novel evidence with two previous ones, namely: (i) the distinct electrophysiological correlates of NPs vs. VPs [4]; (ii) the distinct cortical connectivity related to NPs and VPs [6]. The whole picture provides a first electrophysiological fine-grained contrast of two basic syntactic units, namely NPs and VPs, having excluded the confounding factor of phonological information. We found activations correlating with syntactic and lexical processing in different electrodes, different frequency bands and different time windows. These results showed that both syntactic and lexical information are important for language processing but rely on distinct computations. Moreover, the present study showed that surprisal models based only on morphological information do not play a significant role. Syntactic information is needed to explain brain activations.

Our research on neural syntax processing not only enhances our understanding of language in the brain but also offers promising technological prospects. It could lead to the implementation of communication devices for individuals with language disabilities for whom speech prostheses based on motor cortex activity may be ineffective due to the disruption of the language network (e.g., aphasia), as well as more context-aware virtual assistants, revolutionizing how we interact with linguistic computation in the brain.

## Data and code availability

The data and custom *Python* and *MATLAB* code supporting these findings are available from the corresponding author upon reasonable request.

## Acknowledgements

This work was supported by the Italian Ministry for Universities and Research (INSPECT PRIN 2017JPMW4F) and by the Bertarelli foundation.

## Author contributions

Conceptualization: S F C, A M, and S M; Methodology: A C, C B, F A, M G, F B, C R, S F C, S M, A M, and E R; Experimental setup: C B, F B; Data Collection: A C and C B, Software: A C and C B; Validation: A C, C B, F A, and E R; Formal Analysis: A C and C B; Theoretical syntax contribution: A M; Computational linguistic contribution: R F; Lexical analysis: M G; Resources: S M and E R; Data Curation: A C and C B; Writing - Original draft: A C, C B, and M G; Writing - Review and Editing: all authors; Visualization: A C; Supervision: A M, and E R; Project Administration: S F C, A M, and S M; Funding Acquisition: S M, A M, and E R. C B contributed equally as first author and has the right to put her name first in her CV.

## Competing interests

The authors declare no competing interests.

## Notes

### Competing Interest Statement

The authors have declared no competing interest.

